# Target-distractor competition modulates saccade trajectories in space and object-space

**DOI:** 10.1101/2022.11.03.514759

**Authors:** Caroline Giuricich, Robert J. Green, Heather Jordan, Mazyar Fallah

## Abstract

Saccade planning and execution can be affected by a multitude of factors present in a target selection task. Recent studies have shown that the similarity between a target and nearby distractors affects the curvature of saccade trajectories, due to target-distractor competition. To further understand the nature of this competition, we varied the distance between and the similarity of complex target and distractor objects in a delayed match-to-sample task to examine their effects on saccade trajectories and better understand the underlying neural circuitry. For trials with short saccadic reaction times (SRTs) when target-distractor competition is still active, we found a robust effect of distance consistent with saccade vector averaging, whereas the effect of similarity suggested the existence of an object-based suppressive surround. At longer SRTs there was sufficient time for competition between the objects to complete and the distractor to be inhibited, which resulted in saccade trajectory deviations exhibiting the effects of a spatial suppressive surround. In terms of similarity, as the target-distractor similarity decreased, the initial saccade angle shifted towards the target, reflecting stronger distractor inhibition. There were no interactions between distance and similarity at any point in the time-course of target-distractor competition. Taken together, saccade trajectories reflect target-distractor competition that is affected independently by both spatial and object-space suppressive surrounds. The differences in saccade trajectories at short and long SRTs distinguish between active and completed decision-making processes. Thus, saccade responses are more beneficial than manual responses in studies of decision-making models.

**Significance Statement:** This is the first study to determine that the distance and similarity between visual objects independently affect saccade trajectories driven by the target-distractor competition process. Thus, spatiotemporal and object identity factors separately feed into saccade planning and execution, resulting in modulations of saccade trajectory metrics which are based on spatial and object-space suppressive surround mechanisms. Furthermore, this modulation of trajectory metrics distinguishes between active and complete decision-making processes. The findings are important for understanding the dynamic networks guiding target selection and are relevant for further development of decision-making models, as well as eye-tracking applications in health and disease.

## Introduction

Two potential saccade goals in the environment compete for attention with the saccade made to the target curving towards the interesting distractor (Findlay and Harris, 1984; Van Gisbergen et al., 1987; Minken et al., 1993; Port and Wurtz, 2003; McPeek and Keller, 2001; McPeek et al., 2003). Curvature towards the distractor results from unresolved target-distractor competition and is related to the neural processing of both objects in the superior colliculus and frontal eye fields (McPeek et al., 2003; McPeek, 2006), due to distractor neural activity increasing above baseline (McPeek and Keller, 2001; McPeek et al., 2003; Port and Wurtz, 2003). With enough time between stimulus onset and saccade initiation, target-distractor competition fully resolves, suppressing the distractor neural activity below baseline causing saccades to curve away instead (Doyle and Walker, 2001, 2002; McSorley et al., 2004; White et al., 2012). Thus, saccadic reaction time (SRT) plays a large role during target selection; shorter SRTs result in curvature towards and longer SRTs result in curvature away from the distractor (Theeuwes and Godijn, 2004; Walker et al., 2006; McSorley et al., 2006; Mulckhuyse et al., 2009; Hickey and van Zoest, 2012).

The similarity between competing objects can also affect the saccade curvature in a target selection task. This is an example of behavioural relevance, how bottom-up feature processing and top-down task demands are integrated into a priority map of the visual field. Studies have investigated the effect of similarity on saccade trajectories by varying low level features such as color and orientation; there is more curvature towards a distractor that is color congruent than color incongruent (Ludwig and Gilchrist, 2003; Mulckhuyse et al., 2009). Kehoe et al. (2018a) examined the effects of similarity on saccade curvature for complex objects rather than simple features and found that the more similar the distractor, the less saccade curvature was produced. They suggested the oculomotor system reweights the competing saccade goals, putting the strongest weight on the most behaviourally relevant object, with distractors receiving weights based on their similarity to that target. These weighted vectors are then integrated to produce the final saccade trajectory.

Studies investigating the effect of a distractor on saccade trajectories often keep the distractor at a set distance to the target and do not vary its similarity, treating the distractor as a placeholder for a competing motor plan (Sheliga et al., 1995). To understand the interplay of distance and similarity on saccade trajectories, we simultaneously varied the angular distance in an egocentric reference frame and the similarity between a target and distractor in a delayed match-to-sample task. We predicted that the effects of distance on saccade trajectories would be more complex than a linear relationship due to attentional suppressive surrounds, which have been shown to follow a Difference of Gaussian (DoG) pattern, predicted by the Selective Tuning model of visual attention (Tsotsos, 1990; Tsotsos et al., 1995; Cutzu and Tsotsos, 2003; Hopf et al., 2006). A close distractor falls into the attentional spotlight, centered on the target. As the distractor is placed further away from the target, it is suppressed according to a gradient with the most suppression at a medium distance, caused by pruning connections irrelevant to the stimulus-of-interest (Cutzu and Tsotsos, 2003; Yoo et al., 2018). This suppression is reduced as distance increases until it disappears outside the range of the suppressive surround.

Attentional suppressive surrounds have also been found in feature-space. For example, as the orientation of a distractor shifts further away from that of the target in feature-space, attention follows a suppressive surround pattern (Tombu and Tsotsos, 2008). Feature-based suppressive surrounds have been found for basic features such as color, orientation, and direction of motion (Tsotsos et al., 2005; Tombu and Tsotsos, 2008; Störmer and Alvarez, 2014; Yoo et al. 2018). Here, we investigated if object-based suppressive surrounds exist when attending to higher order, complex objects.

In the present study, we investigated how spatial distance, similarity, and the interplay between them affect saccade programming through examining spatial and object-based suppressive surrounds. We hypothesized that with enough time for target-distractor competition to be resolved, the distractor would be suppressed according to a spatial suppressive surround with a DoG-shaped effect on saccade trajectory deviations. When varying the target-distractor similarity, we expected that if an object-based suppressive surround exists, we would again find a DoG modulation of saccade trajectory deviations, but that this would be limited by distance since similarity computations depend on local competition within a given visual area.

## Materials and Methods

### Participants

Twenty-six participants (18-43 years old, 6 male) took part in this experiment, all with normal or corrected-to-normal vision. Participants were naïve to the purpose of the experiment and received partial course credit for participating, when applicable. They provided written informed consent prior to beginning the experiment. Human subjects were recruited at a location which will be identified if the article is published. [Author University]’s Human Participants Review Committee approved this study.

### Stimuli

The stimuli used were developed in-house using MATLAB (Mathworks, Natick, MA).

They consisted of a combination of 6 or 7 vertical and horizontal line segments (1° × 0.08°) positioned orthogonal to each other. This number of line segments was used to ensure holistic representations of the objects are used rather than depending on human visual working memory capacity, as determined in prior published studies. These stimuli were created to be novel to participants, and thus do not include letters or numbers in the English language. Line segments were white on a black background (white: CIExy [0.29, 0.30], luminance 126.02 cd/m^2^, black: CIExy [0.27, 0.26], luminance 0.20 cd/m^2^) and were put together to fit within a 2° × 2° box. Two sets of stimuli were used and shuffled throughout the experiment (Figure 1).

**Figure 1.**
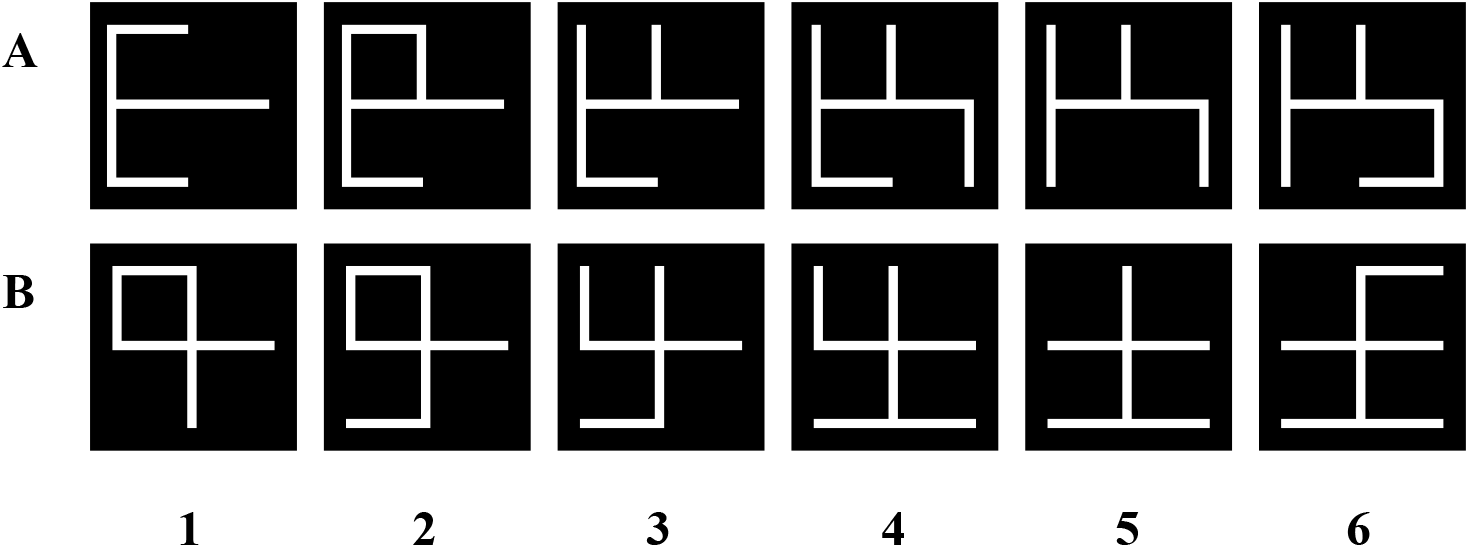
Stimulus sets A and B. The stimuli were numbered from 1-6 (left to right). The difference between stimulus number gives rise to the number of line differences (objective similarity levels). *Apparatus*

Experimental control was maintained with Presentation software (Neurobehavioral Systems, Berkeley, CA). Participants ran the experiment in a dark room on a 21-in CRT monitor (60Hz, 1024×768), 57cm away from a head and chin rest. Eye position was recorded from the left eye using an infrared eyetracker (500Hz; EyeLink II: SR Research, Ontario, Canada). Eye position was calibrated at the start of the experiment and during the experiment, as necessary. Participants responded via a serial response box (Cedrus, San Pedro, CA; 6 participants) or mouse (Dell; 20 participants).

### Procedure

We used a delayed match-to-sample task where participants needed to search for a previously shown target object amongst a distractor object after a short delay from target preview. Participants started each trial when shown a small, white fixation cross (0.4° × 0.4°) at the centre of the screen by pressing a button (serial response box: center button; mouse: left button). The target object was then previewed on the screen (Figure 2A). Once ready, the participants pressed the button again, which replaced the preview with a central fixation cross. After fixating the cross for 200ms, the target and distractor objects appeared simultaneously at isoeccentric points around a circle of radius 8 dva. Participants were instructed to use their peripheral vision to identify the target while maintaining central fixation and respond by moving their gaze to the target. The trial ended when an eye movement was made within a 1.9 dva square box around the target or distractor, or after 750ms if no movement had been made. These time-out trials were reshuffled back into the remaining trial array. An error tone and message were used to indicate incorrect or time-out trials.

**Figure 2.**
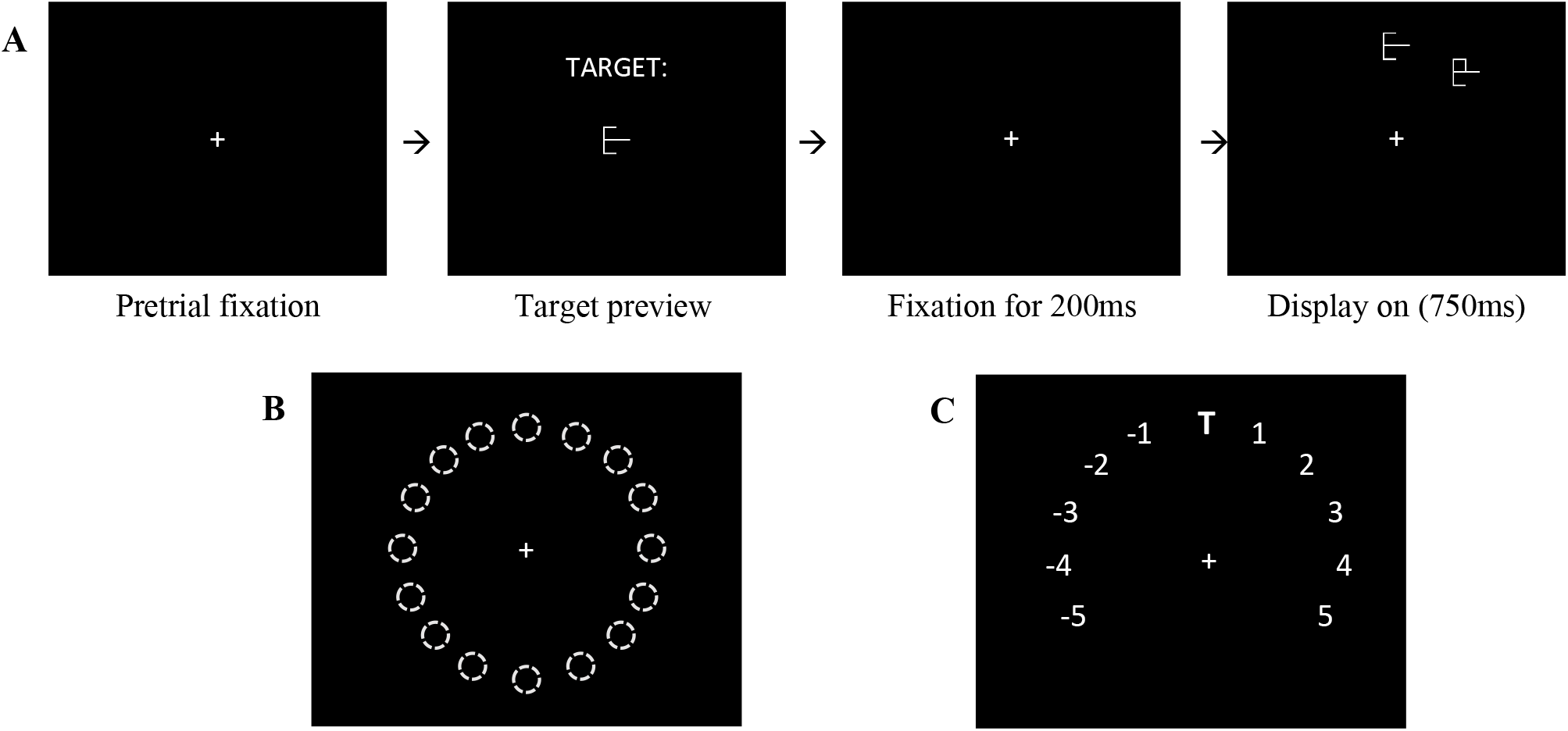
Task paradigm and display layout. ***A***, Task paradigm. Participants fixated on the central cross and pressed a button to move to the target preview. Once ready, they pressed the button again to start the search. Participants stabilized fixation for 200ms then the display came on. The target and distractor remained visible until the participant fixated on an object or until 750ms had passed. ***B***, 16 possible object locations, 22.5° apart, eccentricity of 8 dva. The target was randomly placed at one location and the distractor was placed relative to the target on the clockwise or counterclockwise side. ***C***, AD locations relative to the target. With the target (T) at the top location as an example, the distractor could be placed from 1-5 positions clockwise, or 1-5 positions counterclockwise.

The target and distractor were placed at any of 16 equally spaced positions around the circle (Figure 2B). The target was randomly placed at one location and the distractor was placed relative to the target on either the clockwise or counterclockwise side. Similarity and angular distance between the target and distractor were varied and pseudorandomized for each trial. Lone-target trials without a distractor were considered baseline trials and composed 20% of the total. This resulted in 50 trials per block, with 10 blocks for a total of 500 trials. Participants were given 10 practice trials before block 1 which were not included in analysis. The experiment took ~45mins to complete. All participants included in the analysis completed the full 500 trials.

### Similarity and angular distance

Objective similarity (OS) was determined by the number of line segment differences between two given stimuli, defined by adding or removing individual line segments (Figure 1). Target-similarity differences were confined to the range OS 1-4. Angular distance (AD) was calculated as the absolute difference between target and distractor locations in polar angle (Figure 2B, C). An AD range of ±1-5 locations (spread out by 22.5 deg) was used in this experiment (Figure 2C).

### Analysis

#### Task performance

The behavioural accuracy score is the proportion of trials where the participants correctly made a saccade to the target object. A binomial test was performed on the behavioural accuracy scores for each participant to test that they were more accurate than chance (50%). No participants were excluded from analysis since all scores were significantly better than chance.

#### Saccade detection

In-house written MATLAB scripts were used to filter saccades for each trial. We defined saccades as having a peak velocity over 50°/s, and for at least 8ms, a velocity that exceeded 20°/s. Only correct saccades in which the participants made a saccade to the target were included for analysis. The first saccade of each trial was analyzed and trials in which corrective saccades were needed to reach the target or blinks occurred were excluded from analysis. Saccades with an amplitude < 1° (1.26%) or with an SRT < 100ms (2.96%) were also excluded. In total, 83.82% of correct saccades were included for further analysis.

#### Saccade metric calculations

Five saccade metrics were calculated for each saccade included in the analysis: initial angle, endpoint deviation angle, sum curvature, max curvature, and angle at max curvature. For all saccade metric calculations, the starting position of the saccade was shifted to the origin and rotated so the target position was on the positive y-axis at 8 dva. Initial angle was calculated as the angle between the line from the start of the saccade to the target, i.e. the y-axis, and the point 20% along the saccade trajectory. Endpoint deviation was calculated as the angular difference between the last point of the saccade trajectory and the target, i.e. the y-axis. The curvature metrics were calculated with the saccade rotated so that both the start and end points lay on the positive y-axis. Sum curvature was calculated as the sum of all x-values from the points along the saccade trajectory. Max curvature was the x-coordinate value of the point with the maximum absolute x-value along the saccade trajectory. Angle at max curvature was the angular difference between a straight saccade, i.e. the y-axis, and the max curvature coordinate. These metrics were baseline-subtracted using the average lone-target (distractor-absent) trial metrics at the corresponding target location of that trial. Saccade metrics calculated from trials with negative ADs (counterclockwise placement of distractor relative to target) were reflected and collapsed with the clockwise distractors’ positive AD metrics. This kept the distractor on the positive x side of the y-axis and resulted in 5 total ADs for analysis. All positive metrics signify deviations towards the distractor, and negative metrics signify deviations away from the distractor.

#### Average saccade trajectories

Average saccade trajectories were calculated and plotted using prior published techniques. The x,y coordinates of saccades were separated by OS and/or AD and averaged together by first shifting the trajectory so that the first point was at the origin, and then taking points within 10% sections of the trajectory. Those points were averaged together to create an average saccade to visualize the calculated metrics. We baseline-corrected the x-coordinates of the trajectories by subtracting the values from the average lone-target trial trajectory of the same trial location. Shaded regions around each average trajectory represent one standard error of the mean (SEM).

#### Metric plots and curve fits

Averages of the five metrics were plotted across the four OS levels and five ADs. SEM was calculated and plotted as error bars. We fit each plot with linear, quadratic, sigmoid, Gaussian, Difference of Gaussian (DoG), and exponential curves. We assessed the goodness of fit using the coefficient of determination (R^2^), Akaike information criterion (AIC), and the p-value associated with the F-test of the regression analysis. Our experimental paradigm did not support the use of a DoG curve fit since we did not have the full range of data to represent the excitatory peak of a DoG curve that would occur between our +AD and -AD conditions. Thus, a single Gaussian fit was used to model the suppressive effect over collapsed ADs. Data was also split by OS across ADs to determine the effect of OS at each AD. For each metric, a 5 AD x 4 OS univariate ANOVA was performed in SPSS (v27, IBM) for short and long SRTs separately. Scheffe post-hoc analysis was used to investigate significant effects across conditions.

## Results

### Saccade target onset asynchrony and split saccadic reaction times

Each participant’s behavioural accuracy was significantly above chance, with a mean accuracy across participants of 74%. Only correct trials were included in analysis, where the participants made a saccade to the target. Saccade frequency, trial accuracy, and all five metrics were plotted across averaged saccade target onset asynchrony (STOA) values. STOA was calculated as the time between saccade initiation and target onset, where saccade initiation was taken to be at 0ms. STOA values were averaged over a sliding window of 10ms from −500 to - 100ms. Each point plotted was an average of the metric data from trials with saccadic reaction times (SRTs) that fell within the sliding 10ms window. Trials with SRTs greater than 500ms (2.5% of total trials) were treated as outliers that were not representative of the rest of the data and were not included in analysis.

Plotting saccadic frequency (the number of trials within a given 10ms sliding window in which a saccade was initiated as a proportion of total correct trials) over STOA showed the peak frequency of saccade initiations occurred at ~150ms, as would be expected with visually evoked saccades (Figure 3A). After this peak, there was a general decrease in saccade initiation frequency as STOA increased. Accuracy over STOA (plotted as the percentage of trials that participants made a saccade to the target as a proportion of all total trials) showed that the longer the participants took to process the display, the less likely they were to incorrectly make a saccade to the distractor object, as reflected in the STOA (Figure 3B). There was also more variance in accuracy at long SRTs.

**Figure 3.**
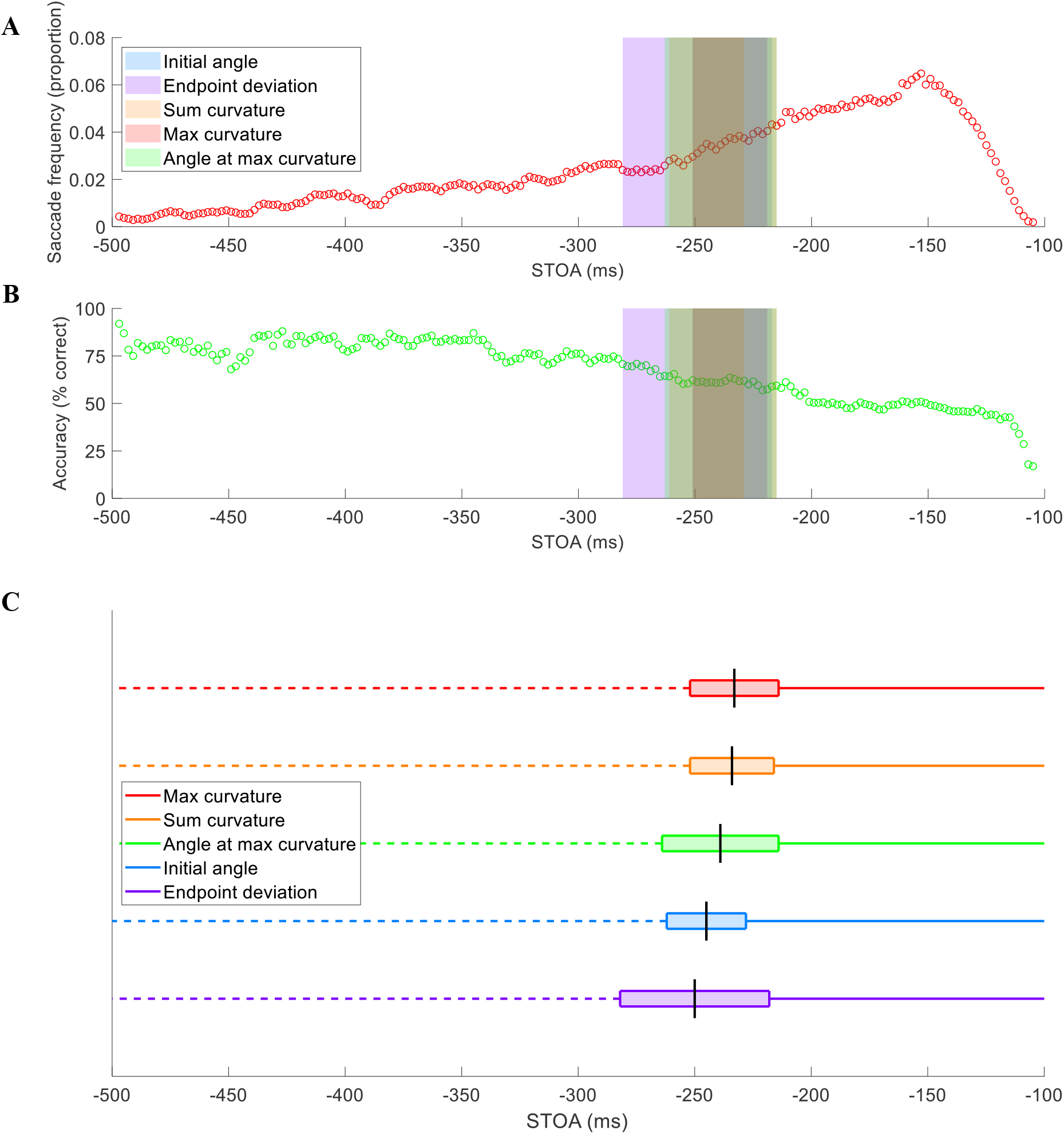
Saccade frequency, accuracy, and metrics over time. Shaded colored regions cover the middle saccadic reaction time (SRT) period (transition period) for each metric that was removed from analysis. ***A***, Saccade frequency plotted over saccade target onset asynchrony (STOA) as a proportion of total trials. ***B***, Accuracy plotted over STOA as a percentage of total trials. ***C***, Transition period ranges by metric over SRT, ordered by midpoint from earliest to latest. The transition period is represented by the box and the midpoint is shown with a black vertical line. Each solid-colored line represents the short SRT period where that metric significantly deviated towards the distractor. Each dashed-colored line represents the long SRT period where that metric significantly deviated away from the distractor.

Figure 3C shows that significant positive deviations (from a straight line vector to target) towards the distractor occurred for short SRTs, and significant negative deviations away from the distractor occurred for long SRTs. The SRT range that fell in the middle was not significantly different than zero for each metric (see Table 1). This ‘transition period’ indicated the range of time where the process flipped from positive to negative deviations. To test if the transition period consisted of proportional mixtures of significant positive and negative deviations or saccades that were not significantly deviated, the metric data from trials with SRTs within the transition periods specific to each metric were plotted as histograms. Each metric histogram fit well with a single Gaussian distribution compared to a dual Gaussian, all with R^2^ > 0.96 and p < 0.001. Therefore, the transition periods reflect the oculomotor system switching from an excitatory to an inhibitory distractor. The saccade metrics were ordered by midpoint of each of their transition periods from earliest to latest (Figure 3C). The curvature metrics reached the midpoint earlier than the angle metrics. All subsequent analyses were split for short and long SRTs outside of the transition periods. It is important to note that trials with short SRTs made up 55% of total trials included in analysis, whereas trials with long SRTs made up 30%. For each analysis split by metric, the inclusive, metric-specific short and long SRT ranges were used, and the transition period was removed from analysis (see Table 1). The maximum spanning short and long SRT metric ranges were chosen as the ranges for the average saccade analysis as this analysis did not require data to be split by metric.

**Table 1.**
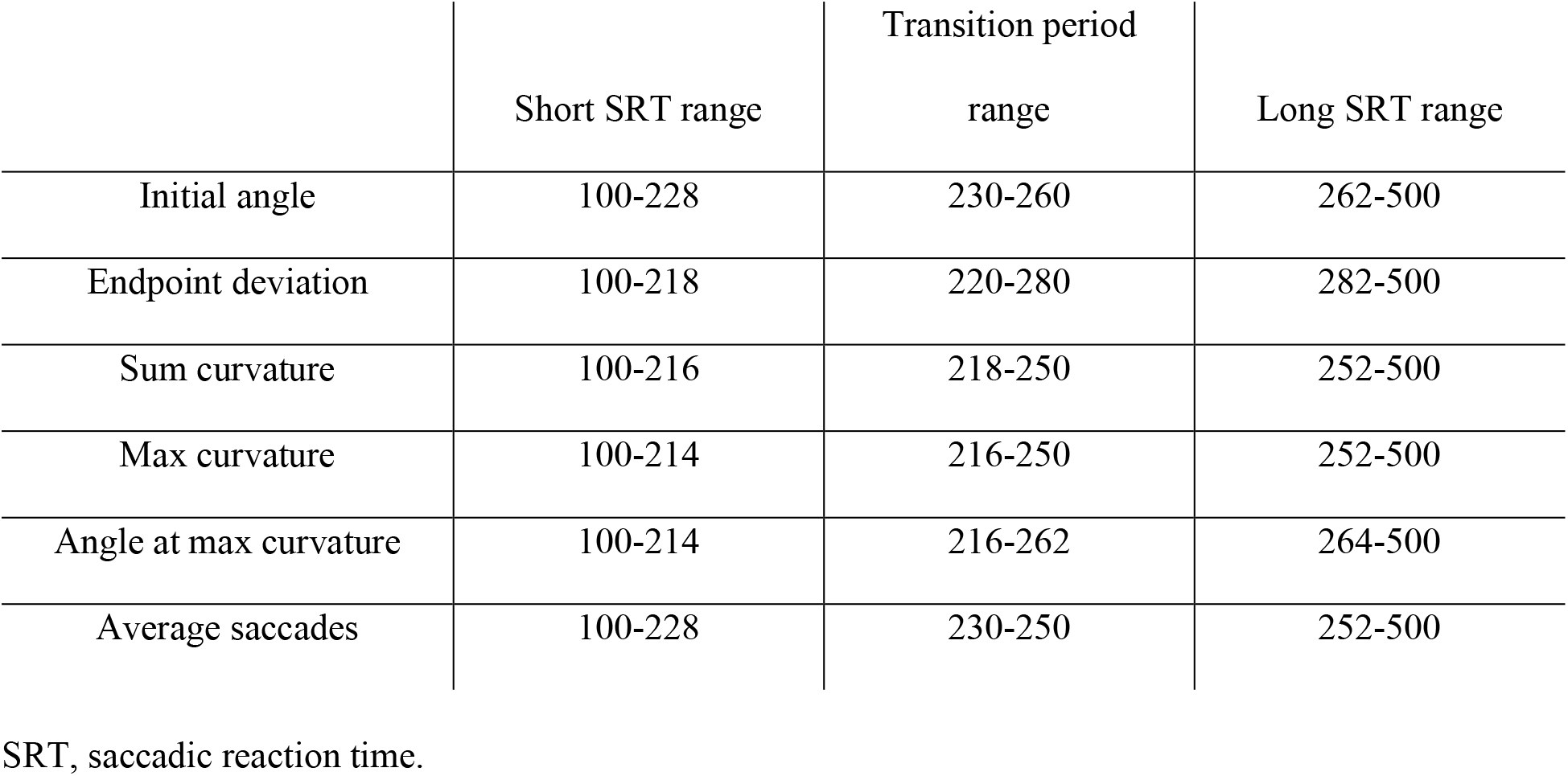
Short SRT ranges, transition period, and long SRT ranges (ms) for each metric and the average saccade analysis.

|  | Short SRT range | Transition period<br>range | Long SRT range |
| --- | --- | --- | --- |
| Initial angle | 100-228 | 230-260 | 262-500 |
| Endpoint deviation | 100-218 | 220-280 | 282-500 |
| Sum curvature | 100-216 | 218-250 | 252-500 |
| Max curvature | 100-214 | 216-250 | 252-500 |
| Angle at max curvature | 100-214 | 216-262 | 264-500 |
| Average saccades | 100-228 | 230-250 | 252-500 |

SRT, saccadic reaction time.

### Average saccades

Average saccade trajectories were plotted across angular distance (AD) and objective similarity (OS) for short and long SRTs (Figure 4). The average saccades show saccades deviated positively for short SRTs versus negatively for long SRTs. When looking at the differences between OS levels and ADs, distance had a greater effect on saccade trajectory than similarity for short SRTs. Figure 4A qualitatively shows a clear distinction between the average trajectories for AD = 1, 2 compared to AD = 3, 4, 5 in the initial angle and amount of curvature towards the distractor. On the other hand, Figure 4B shows that the average trajectories for each OS were clustered together. For long SRTs, the angular distance variations (Figure 4C) showed evidence of a spatial suppressive surround; the average saccade trajectories away from the distractor were more deviated for AD = 2, 3 than for AD = 1, 4, 5. In comparison, the objective similarity differences (Figure 4D) showed that the average saccade trajectories were more spread out between each OS compared to those at short SRTs (Figure 4B).

**Figure 4.**
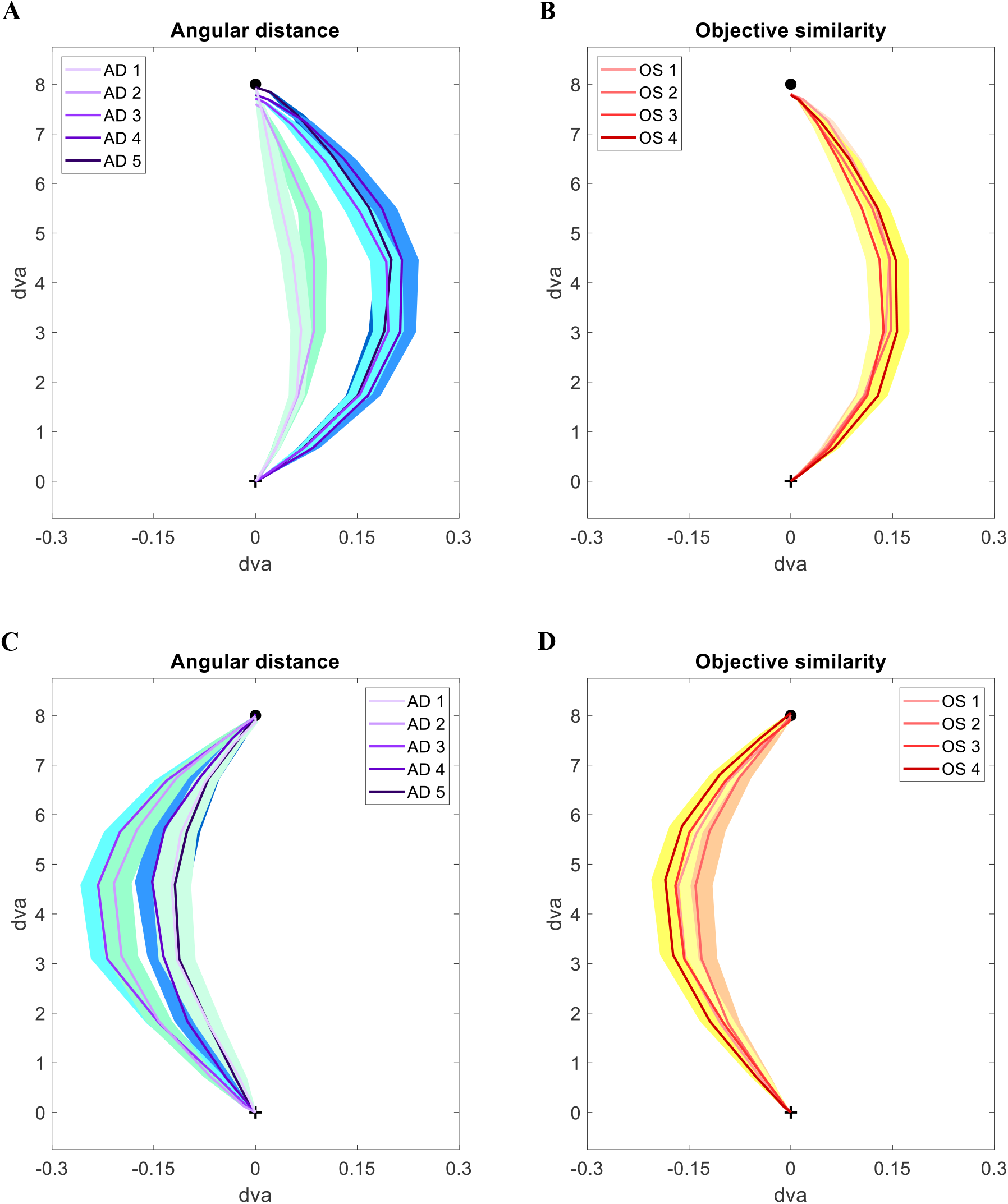
Average saccade plots for short and long SRTs split by angular distance (AD) and objective similarity (OS). Shaded regions represent the SEM for each average point along the trajectories. The black circle represents the target position at 8 dva. ***A***, Average saccades for each AD for short SRTs. ***B***, Average saccades for each OS for short SRTs. ***C***, Average saccades for each AD for long SRTs. ***D***, Average saccades for each OS for long SRTs.

### Angular distance effects

The average metric values were split by short and long SRTs and plotted against AD (Figure 5A, B). Two-way ANOVAs for each metric across short SRTs showed main effects of AD (initial angle p < 0.001, endpoint deviation p < 0.001, sum curvature p < 0.001, max curvature p < 0.001, angle at max curvature p < 0.001) (see Table 2). We next performed Scheffe post-hoc tests on each metric for short SRTs. Initial angle for AD = 1 was significantly more deviated than ADs = 4, 5 (p < 0.001, p = 0.002), and AD = 2 was significantly more deviated than ADs = 4, 5 (p < 0.001, p = 0.003). Endpoint deviation for AD = 1 was significantly more deviated than ADs = 3, 4, 5 (all p < 0.001), AD = 2 was significantly more deviated than ADs = 3, 4, 5 (all p < 0.001), and AD = 3 was significantly more deviated than ADs = 4, 5 (p = 0.030, p = 0.002). Sum curvature for AD = 1 was significantly less deviated than ADs = 3, 4, 5 (all p < 0.001), and AD = 2 was significantly less deviated than AD = 3 (p = 0.008). Max curvature for AD = 1 was significantly less deviated than ADs = 3, 4, 5 (all p < 0.001), and AD = 2 was significantly less deviated than AD = 3 (p = 0.007). Angle at max curvature for AD = 1 was significantly less deviated than ADs = 2, 3, 4, 5 (p = 0.019, rest p < 0.001), and AD = 2 was significantly less deviated than ADs = 3, 5 (p = 0.032, p = 0.008).

**Figure 5.**
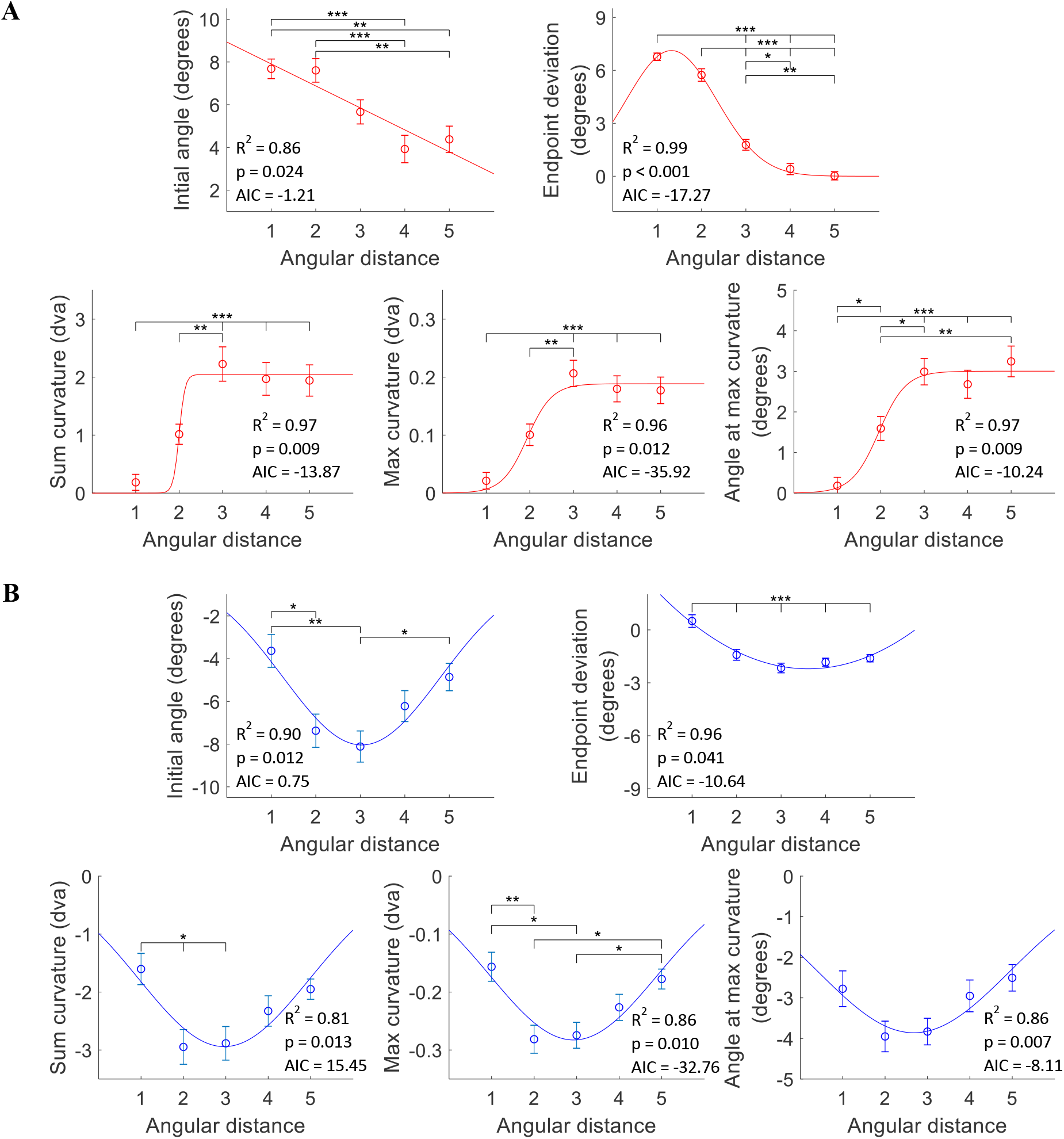
Saccade metric averages over angular distance (AD) with curve fits for short and long saccadic reaction times (SRTs). Curve fits were chosen according to goodness of fit metrics (R^2^, AIC, p). Plots include the respective R^2^, p-values, and AIC. Error bars are SEM. Brackets indicate significant differences (* p < 0.05, ** p < 0.01, *** p < 0.001) from Scheffe post-hoc analysis. ***A***, Metrics over AD for short SRTs. Curve fits from left to right: top: linear, sigmoid; bottom: sigmoid, sigmoid, Gaussian. ***B***, Metrics over AD for long SRTs. Curve fits were all Gaussian except for endpoint deviation which was fit with a quadratic.

**Table 2.** AD and OS main effect ANOVA p-values for each metric for short and long SRTs.

|  | Short SRTs |  | Long SRTs |  |
| --- | --- | --- | --- | --- |
|  | AD | OS | AD | OS |
| Initial angle | < 0.001***† | 0.606 | < 0.001***† | 0.052 |
| Endpoint deviation | < 0.001***† | 0.311 | < 0.001***† | 0.020* |
| Sum curvature | < 0.001***† | 0.746 | 0.001**† | 0.470 |
| Max curvature | < 0.001***† | 0.831 | < 0.001***† | 0.736 |
| Angle at max curvature | < 0.001***† | 0.658 | 0.020* | 0.267 |

p-value significance: \* $p < 0.05$ , \*\* $p < 0.01$ , \*\*\* $p < 0.001$ . † indicates where Scheffe post-hoc analysis showed significant differences between conditions. AD, angular distance; OS, objective similarity; SRT, saccadic reaction time.

For long SRTs, there were main effects of AD on all metrics (initial angle p < 0.001, endpoint deviation p < 0.001, sum curvature p = 0.001, max curvature p < 0.001, angle at max curvature p = 0.020) (See Table 2). Scheffe post-hoc tests for AD vs metrics for long SRTs showed: initial angle for AD = 1 was significantly less deviated than ADs = 2, 3 (p = 0.023, p = 0.002), and AD = 3 was significantly more deviated than AD = 5 (p = 0.025). Endpoint deviation for AD = 1 was significantly less deviated than ADs = 2, 3, 4, 5 (all p < 0.001). Sum curvature for AD = 1 was significantly less deviated than ADs = 2, 3 (p = 0.023, 0.024). Max curvature for AD = 1 was significantly less deviated than ADs = 2, 3 (p = 0.009, p = 0.011), AD = 2 was significantly more deviated than AD = 5 (p = 0.025), and AD = 3 was significantly more deviated than AD = 5 (p = 0.029). There were no significant differences across ADs for angle at max curvature.

We fit each metric plot with linear, quadratic, sigmoid, Gaussian, Difference of Gaussian (DoG), and exponential curves. We used goodness of fit metrics (R^2^, AIC, p-values) to determine the function with the best fit for each metric (see Table 3). The best curve fits for the AD/short SRTs plots were as follows (Figure 5A): initial angle – linear (R^2^ = 0.86, AIC −1.21, p = 0.024); endpoint deviation – Gaussian (R^2^ = 0.99, AIC = −17.27, p < 0.001); sum curvature – sigmoid (R^2^ = 0.97, AIC = −13.87, p = 0.009); max curvature – sigmoid (R^2^ = 0.96, AIC = −35.92, p = 0.012); angle at max curvature – sigmoid (R^2^ = 0.97, AIC = −10.24, p = 0.009). All best fits had significant p-values (F-test from the regression analysis). The best fits for AD/long SRTs (Figure 5B) were as follows: initial angle – Gaussian (R^2^ = 0.90, AIC = 0.75, p = 0.012); endpoint deviation – quadratic (R^2^ = 0.96, AIC −10.64, p = 0.041); sum curvature – Gaussian (R^2^ = 0.81, AIC = 15.45, p = 0.013); max curvature – Gaussian (R^2^ = 0.86, AIC = −32.76, p = 0.010); angle at max curvature – Gaussian (R^2^ = 0.86, AIC = −8.11, p = 0.007). Again, all best fits had significant p-values (F-test from the regression analysis). As Gaussian and quadratic curves fit all metrics similarly for long SRTs, excluding endpoint deviation, we relied on Gaussian curves as they are often found for modelling neural mechanisms.

**Table 3.**
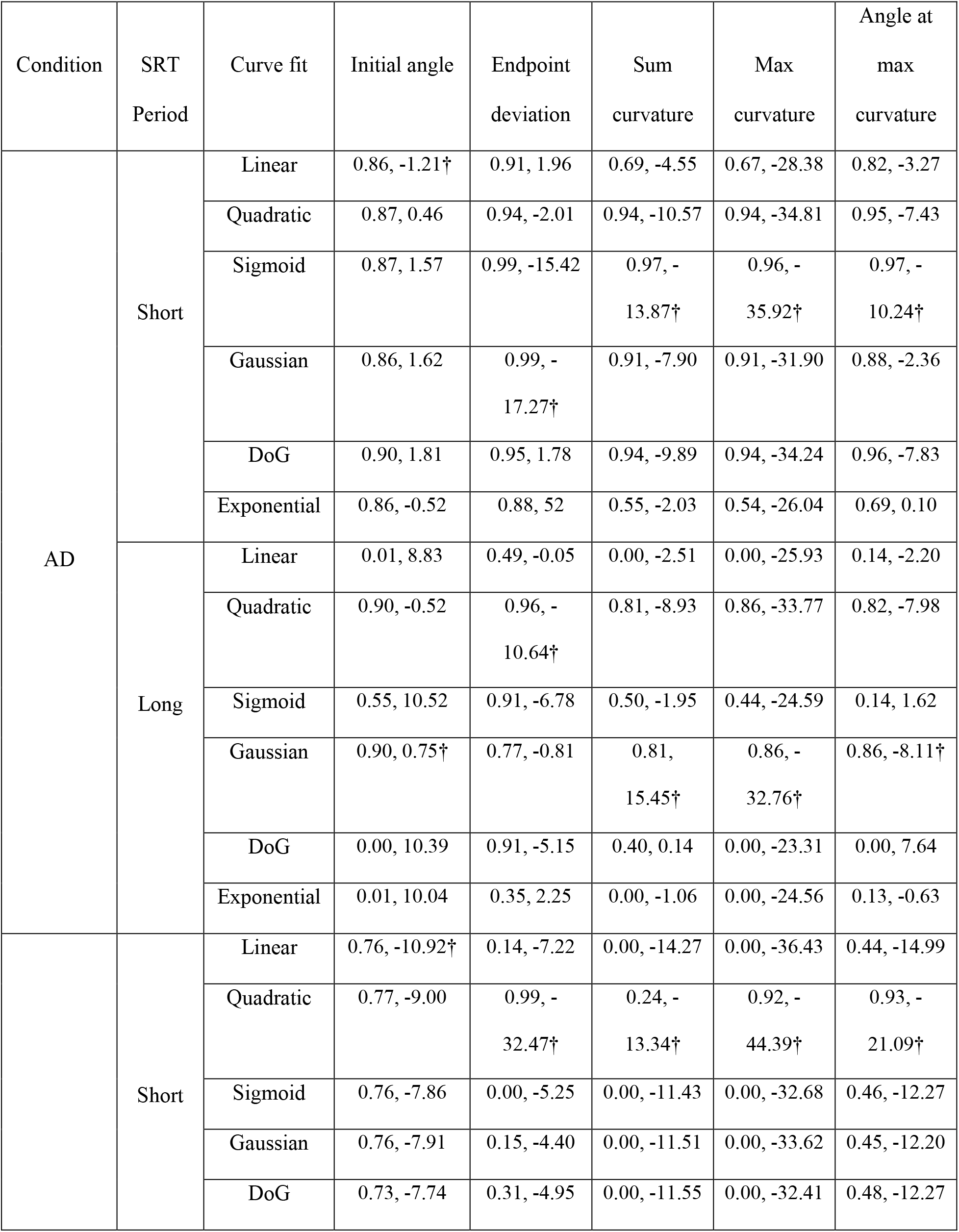
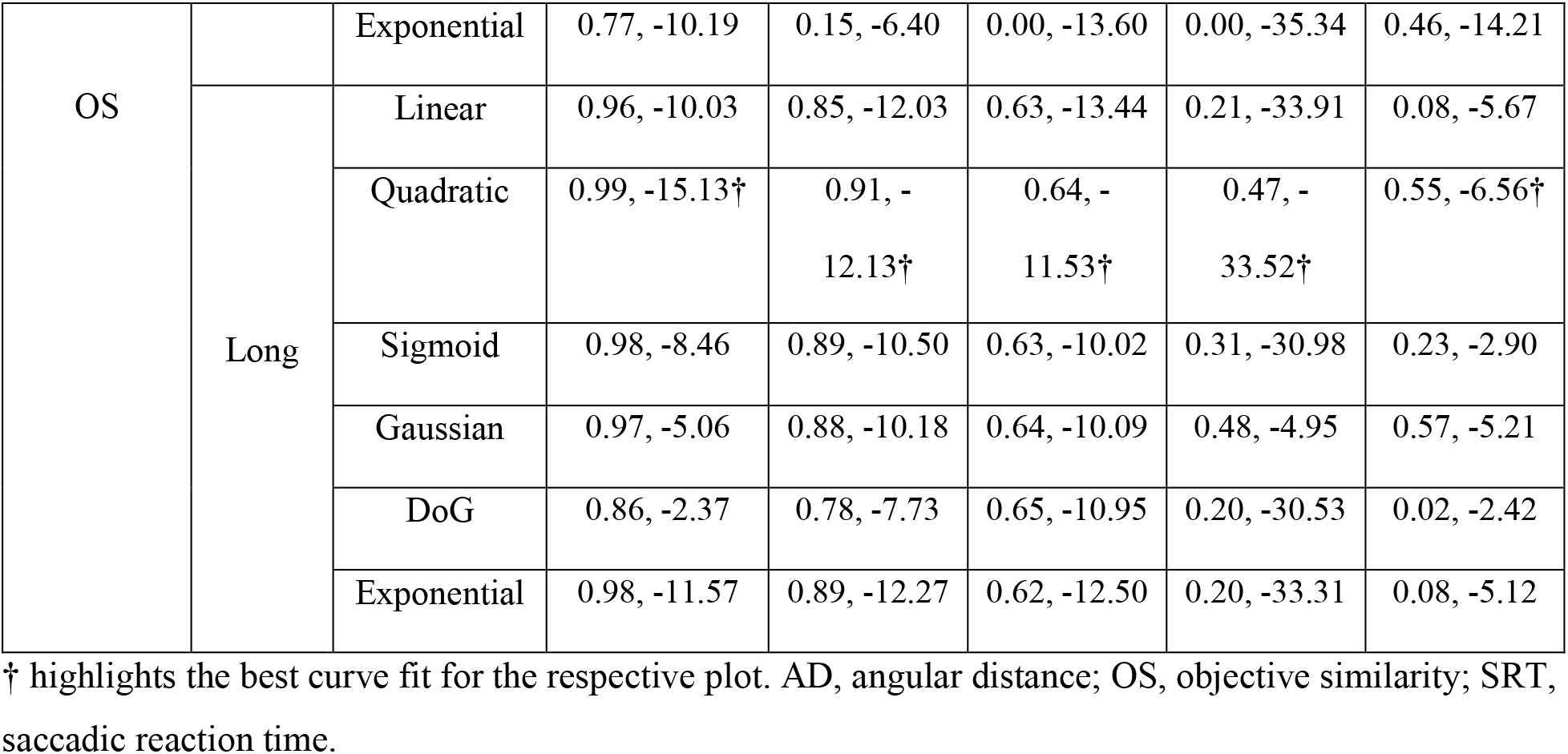
AIC values for each curve fitted to metric plots over AD and OS for short and long SRTs.

The results for the metrics over AD/short SRTs suggest that there was not enough time to inhibit the distractor and resolve the target-distractor competition (Figure 5A), consistent with saccade trajectories shifted towards the distractor. The initial angle and endpoint deviation, reflecting the overall saccade vector, exhibited weighted averaging of the two object locations where distance decreased the weight of the distractor. Curvature measures reflected a different process wherein the strength of the distractor in producing a curved trajectory followed an increase-to-plateau pattern, indicative of an increasing effect of the distractor on saccade curvature that maxes out by 67.5 deg (AD = 3). For long SRTs, target-distractor competition was completed resulting in saccade trajectories shifted away from the distractor. Varying AD resulted in DoG-shaped curves for all metrics, indicative of a spatial suppressive surround where there was the most suppression of the distractor at 67.5 deg (AD = 3, Figure 5B).

### Similarity effects

The average metric values were analyzed against OS for short and long SRTs (Figure 6A, B). For short SRTs, there were no main effects of OS on the metrics. For long SRTs, there was a main effect of OS on endpoint deviation (p = 0.020) (see Table 2), but there were no significant differences across OS levels according to Scheffe post-hoc analysis. There were no significant interaction effects between OS and AD for short or long SRTs.

**Figure 6.**
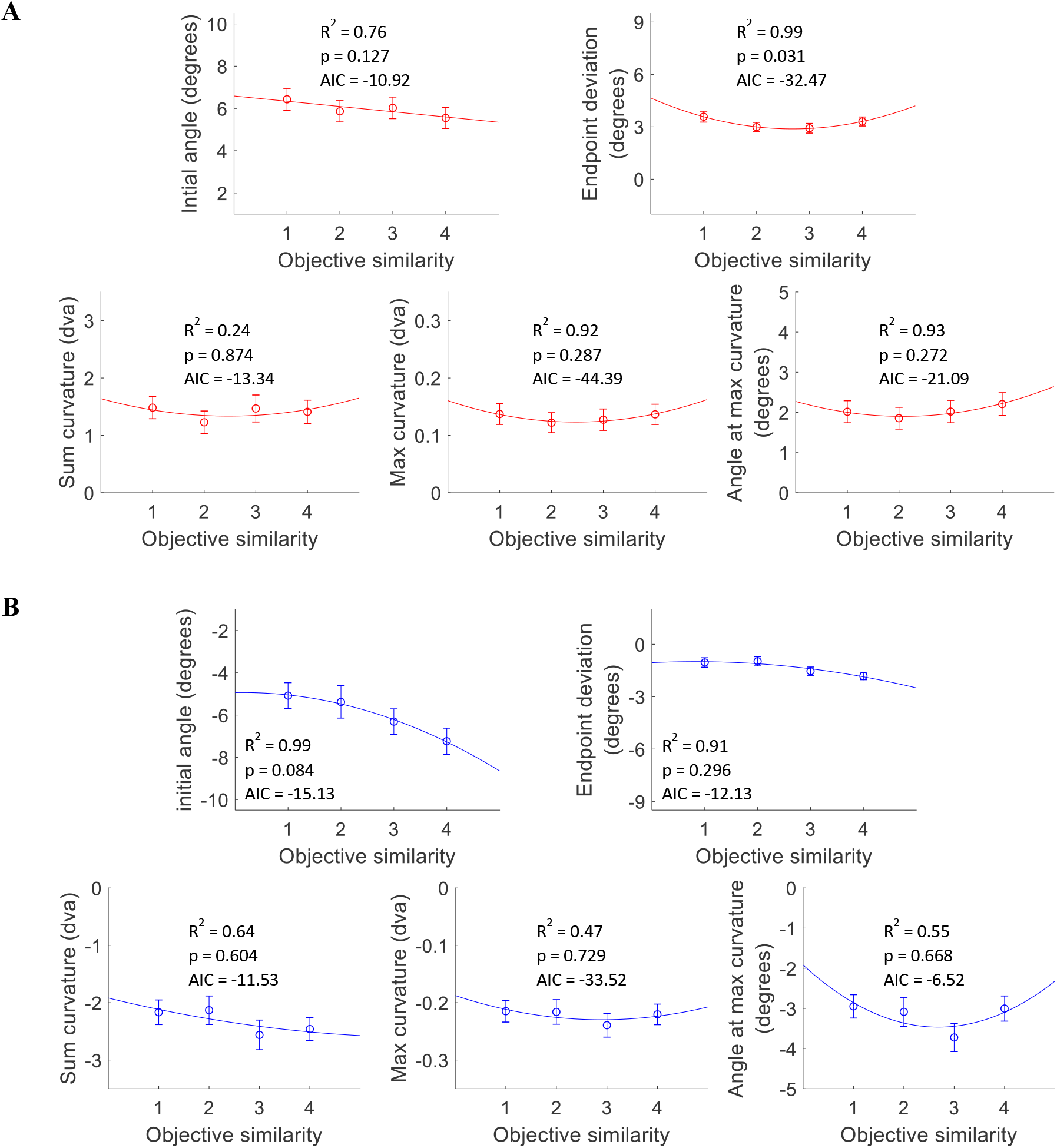
Saccade metric averages over objective similarity (OS) with curve fits for short and long saccadic reaction times (SRTs). Curve fits were chosen according to goodness of fit metrics (R^2^, AIC, p). Plots include the respective R^2^, p-values, and AIC. Error bars are SEM. ***A***, Metrics over OS for short SRTs. Curve fits were all quadratic except linear for initial angle. ***B***, Metrics over OS for long SRTs. Curve fits were all quadratic.

We again fit a range of functions to the metrics plotted against OS. The best curve fits according to goodness of fit measures for OS/short SRTs (Figure 6A) were as follows: initial angle – linear (R^2^ = 0.76, AIC = −10.92, p = 0.127); endpoint deviation – quadratic (R^2^ = 0.99, AIC = −32.47, p = 0.031); sum curvature – quadratic (R^2^ = 0.24, AIC = −13.34, p = 0.874); max curvature – quadratic (R^2^ = 0.92, AIC = −44.39, p = 0.287); angle at max curvature – quadratic (R^2^ = 0.93, AIC = −21.09, p = 0.272). Only the quadratic fit for endpoint deviation was significant. For OS/long SRTs (Figure 6B): initial angle – quadratic (R^2^ = 0.99, AIC = −15.13, p = 0.084); endpoint deviation – quadratic (R^2^ = 0.91, AIC = −12.13, p = 0.296); sum curvature – quadratic (R^2^ = 0.64, AIC = 11.53, p = 0.604); max curvature – quadratic (R^2^ = 0.47, AIC = – 33,52, p = 0.729); angle at max curvature – quadratic (R^2^ = 0.55, AIC = −6.56, p = 0.668).

For OS/short SRTs, there were no significant effects of similarity on saccade trajectories, except for endpoint deviation (Figure 6A). The lack of effect of similarity was expected since at short SRTs there was not enough time to fully cortically process both objects, and inhibit the distractor, to resolve the target-distractor competition. The pattern found for endpoint deviation suggests that an object-based suppressive surround modulates endpoint deviation while discriminating the objects. For OS/long SRTs, we saw a downwards sloping significant trend for initial angle (p = 0.084), showing that the initial saccade vectors angled away from the distractor more when the objects were less similar to each other (Figure 6B).

### Similarity-angular distance split plots

To better investigate the relationship between OS level and AD, we plotted the average metric values across AD with separate lines for each OS level (Figure 7). The overall patterns match those of the averaged metric plots from Figure 5A, B. We fit each line with the best curve fit from the prior AD analysis for their respective SRT periods. For both short and long SRTs, the best fit of the average metric plots (see Table 4) matched well to the individual similarity lines with most curves showing significant fits (p < 0.05) or trends (p < 0.1).

**Figure 7.**
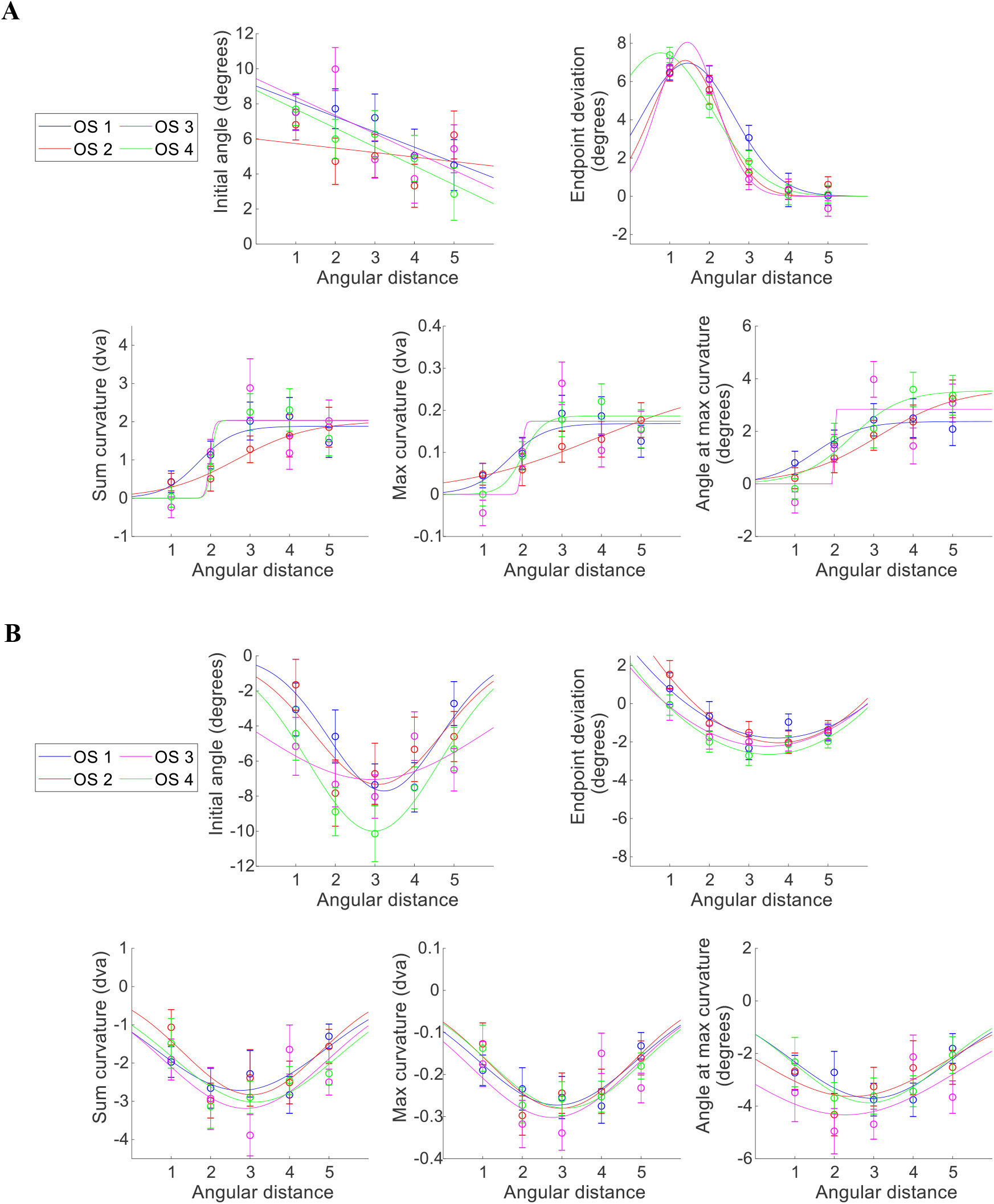
Metrics over angular distance (AD) with separate lines for objective similarity (OS) 1-4, plotted for short and long saccadic reaction times (SRTs). Each plot was fit with the respective best fit per metric from the average metric plots. ***A***, Metrics over AD with split OS lines for short SRTs. ***B***, Metrics over AD with split OS lines for long SRTs.

**Table 4.** R^2^, AIC, p-values for each curve fitted to metric plots over AD separated by OS for short and long SRTs.

| SRT<br>Period | Metric<br>(Curve fit) | OS 1 | OS 2 | OS 3 | OS 4 |
| --- | --- | --- | --- | --- | --- |
| Short | Initial angle<br>(Linear) | 0.83, -2.01, 0.030* | 0.09, 5.49, 0.626 | 0.44, 9.04, 0.220 | 0.89, -2.44, 0.459 |
|  | Endpoint<br>deviation<br>(Gaussian) | 0.99, 16.76,<br><0.001* | 0.987, 16.35,<br>0.009* | 0.99, 17.96, 0.009* | 0.99, 17.73, 0.002* |
|  | Sum curvature<br>(Sigmoid) | 0.83, -5.92, 0.041* | 0.97, -16.50,<br>0.009* | 0.72, 4.04, 0.031* | 0.91, -1.13, 0.004* |
|  | Max curvature<br>(Sigmoid) | 0.77, -28.85,<br>0.054* | 0.97, -41.14,<br>0.007* | 0.69, -22.23,<br>0.047* | 0.93, -32.28,<br>0.028* |
|  | Angle at max<br>curvature<br>(Sigmoid) | 0.90, -8.91, 0.015* | 0.98, -12.29,<br>0.008* | 0.71, 8.07, 0.046* | 0.91, -2.20, 0.038* |
| Long | Initial angle<br>(Gaussian) | 0.88, 18.55, 0.026* | 0.62, 9.47, 0.078* | 0.20, 8.21, 0.048* | 0.94, 0.57, 0.007* |
|  | Endpoint<br>deviation<br>(Quadratic) | 0.76, -0.92, 0.240 | 0.97, -9.15, 0.031* | 0.96, -13.45,<br>0.037* | 0.92, -7.74, 0.080* |
|  | Sum curvature<br>(Gaussian) | 0.63, -4.22, 0.031* | 0.70, -2.89, 0.043* | 0.32, 2.54, 0.087* | 0.67, -4.06, 0.026* |
|  | Max curvature<br>(Gaussian) | 0.77, -30.34,<br>0.018* | 0.73, -27.56,<br>0.031* | 0.34, -21.21,<br>0.085* | 0.85, -32.20,<br>0.012* |
|  | Angle at max<br>curvature<br>(Gaussian) | 0.64, -1.10, 0.031* | 0.51, -0.35, 0.035* | 0.23, 5.50, 0.074* | 0.94, -10.63,<br>0.004* |

\* highlights significant effects ( $p < 0.05$ ) or significant trends ( $p < 0.1$ ). AD, angular distance; OS, objective similarity; SRT, saccadic reaction time.

### Gradient plots

We depicted the significant effects and trends from the average metric plots (Figures 5 and 6) with radial gradient plots in distance (Figure 8) and object-space (Figure 9). The darkest red and blue colors represent the most deviation towards and away from the distractor, respectively. The lighter the color, the closer the metric was to no deviation. Figure 8A depicts the effects of AD on saccade metrics at short SRTs shifted towards the distractor, where initial angle and endpoint deviation showed a decreasing trend as distance increases, whereas the curvature metrics showed an increasing trend to a plateau at AD = 3. Figure 8B illustrates the effects of AD at long SRTs and showed a suppressive surround pattern for all metrics shifting away from the distractor, consistent with a DoG-shaped spatial suppressive zone. Figure 9A depicts the effects of OS on endpoint deviation for short SRTs as a suppressive surround in object-space where there was less endpoint deviation towards the distractor at the middle similarity levels (OS = 2, 3). Figure 9B shows the effects of OS at long SRTs through a decreasing trend for initial angle, further shifting away from the distractor as OS increased.

**Figure 8.**
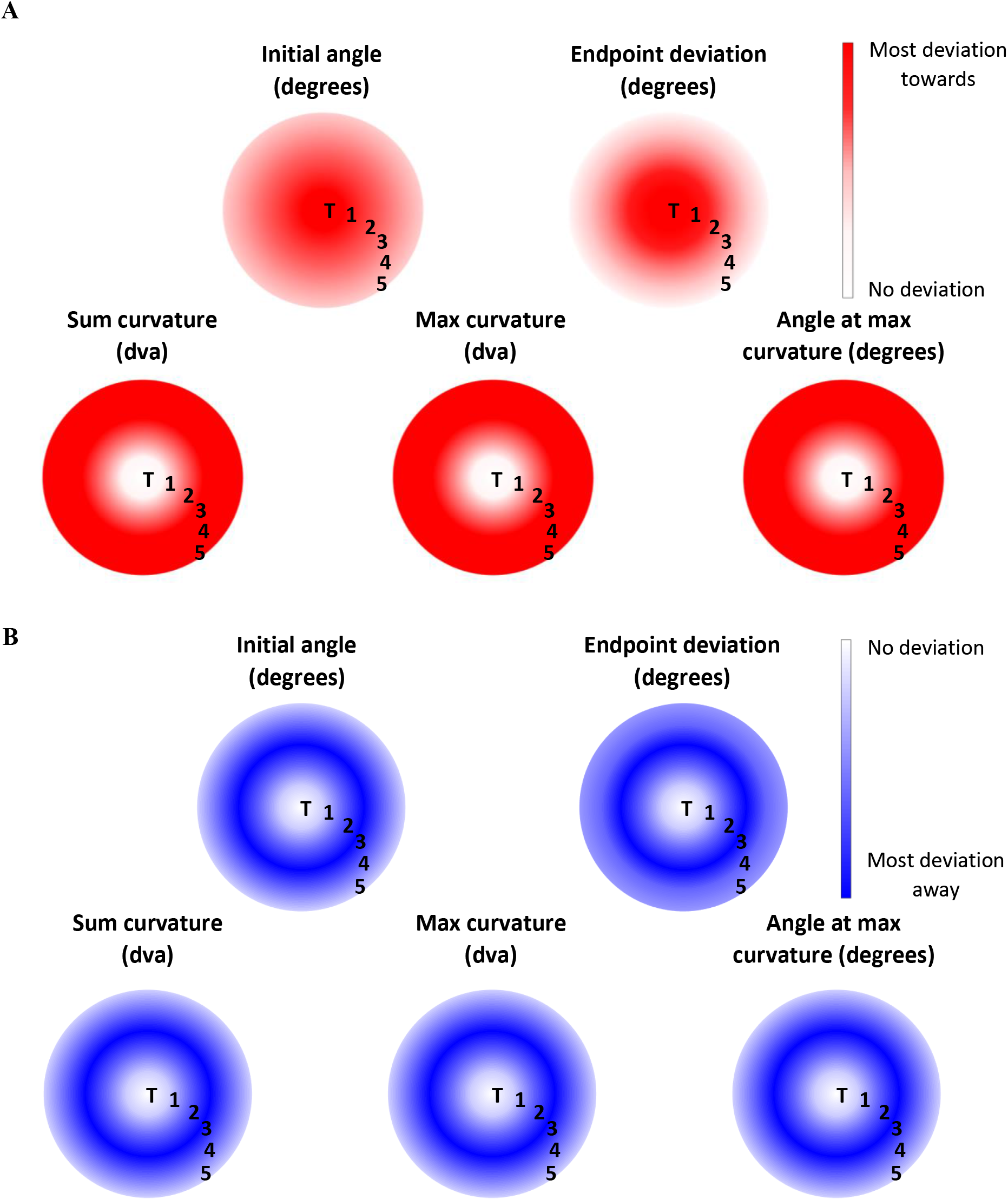
Metric plots depicted as radial gradients over angular distance (AD) for short and long saccadic reaction times (SRTs). Target and distractor positions in space are shown with T and AD 1-5. Darker colors represent the most deviation towards or away from the distractor. White represents no deviation. ***A***, Gradient plots over AD for short SRTs. ***B***, Gradient plots over AD for long SRTs.

**Figure 9.**
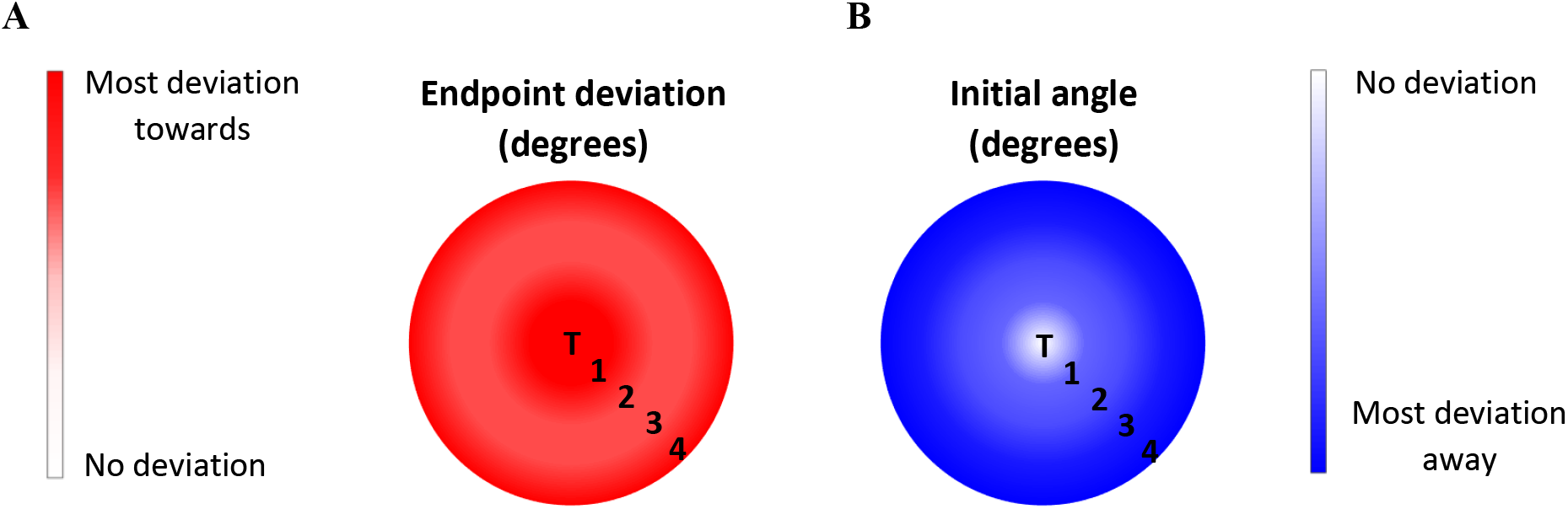
Metric plots depicted as radial gradients over objective similarity (OS) for short and long saccadic reaction times (SRTs). These depictions were created for significant trends (p < 0.1) which occurred for endpoint deviation and initial angle for OS/short SRTs and OS/long SRTs, respectively. Target and distractor positions in object-space are shown with T and OS = 1-4. Darker colors represent the most deviation towards or away from the distractor. White represents no deviation. ***A***, Gradient plot over OS for short SRTs. ***B***, Gradient plot over OS for long SRTs.

## Discussion

We examined the effects of varying the angular distance (AD) and objective similarity (OS) of complex target and distractor objects on saccade trajectories made to the target during a delayed match-to-sample task to elucidate how perceptual and spatial relationships are encoded by the oculomotor system. The time between display onset and saccade initiation influenced saccade direction, prompting us to further analyze the effects of angular distance and similarity based upon saccadic reaction time (SRT). While some prior studies have shown differential effects of distractor attraction and inhibition based on broad timing differences (Theeuwes and Godijn, 2004; Walker et al., 2006; McSorley et al., 2006; Mulckhuyse et al., 2009; Hickey and van Zoest, 2012), we determined the time course of the switch from distractor attraction to inhibition for each of the five saccade trajectory metrics. The effects of SRT on the saccade metrics reflect the development of the competition process between saccade goals; more time between target-distractor onset and saccade initiation resulted in more time to resolve competition and inhibit the distractor. When there was not enough time to resolve the target-distractor competition, there was an effect of angular distance on saccade trajectories, consistent with saccade vector averaging and incomplete winner-take-all (Port and Wurtz, 2003). With increasing SRT, distance modulated the saccade metrics via a spatial suppressive surround. Varying target-distractor similarity had limited effects, only influencing endpoint deviations at short SRTs that reflected an object-based suppressive surround during the active discrimination phase of competition. At long SRTs, varying similarity only affected initial angle, showing saccades increasingly shifted linearly away from the distractor as object similarity decreased.

### Effects of saccadic reaction time on saccade direction

Although we did not purposefully manipulate the SRT, the time taken to decide which object was the target and initiate a saccade naturally varied across trials due to the difficulty of discriminating complex objects. When participants took little time (short SRTs), saccades curved towards the distractor; when participants took more time (long SRTs), saccades curved away from the distractor. This is dependent on the timing of the target-distractor competition process, the time it takes select a target and inhibit the distractor, ~200-220ms (McSorley et al., 2006). With a shorter SRT period (~100-220ms), the target-distractor competition remained unresolved at saccade initiation, resulting in curvature towards the distractor that reflects weighted vector averaging between the target and distractor. With a longer SRT period (~260-500ms), there was enough time to inhibit the distractor causing saccades to curve away from the distractor. In the time between the short and long SRTs (~220ms-260ms), the distractor was in a ‘transition period’ between competing with the target and being inhibited.

However, the transition period ranges varied between each saccade metric. We ordered these ranges from earliest to latest by the midpoint of the transition period, an approximate time for when the metrics flipped from deviating towards to away. The curvature metrics (sum curvature, max curvature, and angle at max curvature) reached the midpoint earlier than the angle metrics (initial angle and endpoint deviation) suggesting that after competition resolves and the process flips to distractor inhibition, saccade curvature is affected sooner than initial angle and endpoint deviation. This is likely because the curvature metrics reflect the residual effects of an incomplete winner-take-all process. Initial angle and endpoint deviation occurred later because they are reflective of weighted vector averaging in saccade planning, based on that prior winner-take-all process. These findings for saccade angle and curvature are consistent with prior studies that found the initial speed and direction of smooth pursuit to a moving target against a moving distractor reflected weighted vector averaging, until the winner-take-all process completed (>300 ms) (Recanzone and Wurtz, 1999; Lisberger and Ferrera, 1997; Ferrera and Lisberger, 1995, 1997). Then, the distractor was fully inhibited, and the target was pursued accurately.

### Effects of angular distance on saccade trajectories

For short SRTs, initial angle and endpoint deviation decreased with increasing angular distance, reflective of weighted vector averaging of saccade plans prior to movement initiation. The saccade angle was shifted more towards the target with increasing distance, suggesting that when planning a saccade to one location, competing plans have decreased strength the further away they are to the saccade goal. As the same pattern was seen for both initial angle and endpoint deviation, they must both be derived from the underlying saccade vector, separate from the curvature metrics in terms of saccade motor planning. Our results further support a prior proposal that the initial direction and endpoint deviation of a saccade are correlated (Van der Stigchel et al., 2007), refuting other models that propose endpoint deviation is independent of the on-line correction (curvature) guiding the saccade back towards the target (McSorley et al., 2004).

The curvature metrics showed evidence of an incomplete winter-take-all process between the locations of the target and distractor, developed from saccade goal competition. As angular distance increased, saccade curvature increased and then plateaued. When the distractor is next to the target, there is little to no competition; the competition increases as the distance between target and distractor increases, until a maximum saccade curvature is reached. This maximum is maintained up through 112.5° (AD = 5), the largest distance in our study. Beyond that range, it is likely to decrease again, e.g. when approaching 180° of separation (opposite vectors).

Overall, these results showed the effects of varying angular distance on saccade trajectories when the target-distractor competition had not had enough time to resolve, since deviations towards the distractor were exclusively seen. Incomplete winner-take-all competition between saccade goals led to saccade curvature that increased with distance until a maximum was reached. With slightly later timing, initial and endpoint deviation angles reflected a weighted vector average for saccade motor plans to target and distractor locations.

For long SRTs, initial angle and endpoint deviation shifted away from the distractor in a Difference-of-Gaussian (DoG) manner as distance increased. The curvature metrics exhibited the same DoG-shaped effects for increasing distance. For both, close distractors shifted saccade plans, but the effects of the distractor decreased through intermediate distances and then re-activated at farther distances. These findings provide evidence of a spatial suppressive surround in oculomotor planning as was predicted and found in visual processing by the Selective Tuning model of attention (Cutzu and Tsotsos, 2003; Hopf et al., 2006; Kehoe et al., 2018b; Yoo et al., 2022). When the distractor was close to the target (AD = 1, 2) it was in the attentional window of the target and less suppression was induced. At the medium distance from target (AD = 3), the distractor exhibited peak suppression from the suppressive surround gradient. Finally, as the distractor moved outside of the suppressive zone surrounding the attended target (AD = 4, 5), suppression was weaker, and curvature similarly lessened. While spatial suppressive surrounds have been found in early and intermediate visual areas (Müller and Kleinschmidt, 2004; Hopf et al., 2006), our results extend this to saccade trajectories derived from the spatial locations of complex objects. Therefore, oculomotor planning is dependent on an attentional priority map with weighted representations of target and distractor locations.

### Effects of similarity on saccade trajectories

We split our similarity results by short and long SRTs to assess how similarity affects the competition phase separately from the distractor inhibition phase after competition resolved. For short SRTs, there were no statistically significant differences between similarity levels when examining the angle and curvature metrics over increasing OS (as the objects became less similar). However, there was a significant DoG-shaped pattern for endpoint deviation, consistent with a non-spatial suppressive surround, found for simple features such as color, orientation, and direction of motion (Tsotsos et al., 2005; Tombu and Tsotsos, 2008; Störmer and Alvarez, 2014; Yoo et al. 2018; Kehoe et al., 2018b). Our results show, for the first time, an object-based suppressive surround active during the discrimination phase of target-distractor competition. When searching for the target, a suppressive surround inhibits the distractor in object-space when it is slightly different from the target. Therefore, the distractor affects endpoint deviation less at these similarity levels, causing saccades to land closer to the target. The object-space suppressive surround is likely mediated by the inferotemporal (IT) cortex, a late stage of ventral stream visual processing that provides object representations (Baylis and Rolls, 1987; Miller et al. 1993; Tanaka, 2003; Gross, 1992), whereas the superior colliculus and frontal eye fields are only capable of discriminating simple features (Kehoe et al., 2018a). Further research is needed to better understand object-space suppressive surrounds.

For long SRTs, the saccade metrics deviated away from the distractor, reflecting distractor inhibition after complete discrimination. As the target and distractor became less similar, the initial angle deviated further away from the distractor. This linear trend reflects increased suppression as an object is more different from its competitor (Perry and Fallah, 2014). There were no effects of similarity on saccade curvature during this late phase of competition. As well, the lack of a DoG-shaped effect from any metric at long SRTs suggests that object identity only mattered during the discrimination phase of target-distractor competition. Once the distractor is fully inhibited, and the competition process is complete, the linear effect of similarity on saccade angle is simply reflected in the initial saccade vector.

A prior study tested the effects of similarity on saccade curvature with a target and two distractors placed 45° from each other. They found that as the similarity between the target and most similar distractor object decreased (as OS increased), saccade curvature (sum and max curvature) and endpoint deviation increased linearly. To compare the present study with the prior study, we analyzed the saccade metrics across similarity for the subset of trials with the distractor at 45° (AD = 2) using linear regressions. For short SRTs, there were no significant linear patterns. For long SRTs, mean endpoint deviation significantly increased as similarity decreased (R^2^ = 0.97, p = 0.02). Initial angle and sum curvature followed the same linear relationship, although not statistically significant (R^2^ = 0.76, p = 0.13; R^2^ = 0.80, p = 0.11). The lack of statistical significance in linear regressions over initial angle and sum curvature was likely due to a lack of statistical power. Since we varied angular distance throughout the experiment, conditions with AD = 2 occurred about 1/5^th^ as many times as in the prior study. Therefore, our findings at 45° are qualitatively consistent with the prior study after the discrimination phase of target-distractor competition has completed and the distractor is inhibited.

### Interaction of angular distance and similarity

While we hypothesized that the effects of similarity on saccade trajectories would be modulated by distance through a multiplicative gain mechanism (Salinas and Abbott, 1997; Connor et al., 1997; Treue and Martínez-Trujillo, 1999; McAdams and Maunsell, 1999; Salinas and Their, 2000), we found no such significant interaction. This suggests that spatial layout and object similarity are independently processed and incorporated into oculomotor planning. When target-distractor competition was active (short SRTs), distance effects were produced by weighted saccade averaging and incomplete winner-take-all, whereas similarity effects on endpoint deviation were consistent with object-space suppressive surrounds. Once the target-distractor competition process has been fully resolved, a spatial suppressive surround is activated resulting in the effects of distance on saccade trajectories. Thus, the stage of the target-distractor competition process gives rise to independent sequential suppressive surround effects first through similarity then distance.

It is important to note that the effects of distance on saccade trajectories were more robust than those of similarity. Angular distance affected all saccade metrics whereas similarity mainly affected initial and endpoint deviation angle, and the effects of distance were greater in magnitude compared to similarity (see Figures 5, 6 and Table 2). Smaller similarity effects on saccade trajectories were likely due to the complexity of the objects and the time given to complete the task. Since the objects were novel shapes, instead of familiar objects, discriminating the target and distractor within the allotted time was difficult. In a task with less emphasis placed on moving quickly and more familiar objects, similarity effects would likely be more robust.

### Oculomotor circuitry and target selection

This study demonstrates that varying the distance and similarity of complex objects, as well as the observed variability in timing of the competition process, affect target selection and saccade planning differently. Saccades initiated during the early stages of target-distractor competition (short SRTs) exhibit weighted vector averaging caused by enhanced neural activity of oculomotor neuron populations encoding the competing saccade goals (Port and Wurtz, 2003; Godijn and Theeuwes, 2002; Findlay and Walker, 1999). The final motor plan is a result of reweighting potential saccade goals based on their behavioural relevance and priority, which for this task is the similarity of the objects in question to the previewed target (Kehoe et al., 2018a). On-line corrections (evidenced by saccade curvatures) pull the overall saccade trajectory back towards the target since its initial vector is more towards the distractor due to the weighted average of the object locations. With more time (long SRTs), the target-distractor competition resolves, consistent with the population of neurons encoding the unchosen saccade goal decreasing in activity (McPeek et al., 2003; McPeek, 2006; White et al., 2012), producing curvatures away from the distractor. Spatial and object-space suppressive surrounds develop around the attended target, further modulating the effect of the distractor on the saccade vector.

The spatial layout of objects in the visual field is encoded in the oculomotor system through attentional priority maps where neurons encoding objects of interest (through both bottom-up and top-down features) have increased activity (Fecteau and Munoz, 2006; White et al., 2017; Fernandes et al., 2014). With sufficient time, the suppressive zone surrounding the attended target develops in the oculomotor system. Studies have shown a link between spatial attention and oculomotor areas (Moore and Fallah, 2001, 2003; Moore et al., 2003; Krauzlis et al., 2013; Bisley and Goldberg, 2003), which suggests that spatial surround suppression developed in visual processing areas feeds into attentional priority maps in the oculomotor system.

In contrast, the target-distractor similarity in this delayed match-to-sample task only affected saccade trajectories while the discrimination process was active. Complex visual information is processed in later regions of the ventral stream, such as areas TE and TEO (Baylis and Rolls, 1987; Miller et al. 1993; Tanaka, 2003). While our target and distractor are processed and discriminated in these higher order visual regions, our results show that their output projects to the oculomotor system for saccade planning and distractor inhibition. Once the discrimination process completes, similarity no longer affects saccade planning. Therefore, spatial layout and object similarity are processed in their respective visual processing areas, with distance and similarity information independently feeding into different aspects of oculomotor planning.

### Applications to decision-making models

Decision-making models define the functions of the active decision-making process and exit points at which the decision is made. These two stages, active and complete, were clearly distinguishable from their effects on saccade trajectories. Our results show that there are two distinct stages in which saccades can be made. When the target-distractor discrimination process is active, as evidenced by short SRTs, saccades curve towards the distractor. When it is complete, with long SRTs, saccades curve away from the distractor. The range of SRTs and resultant effects on saccade trajectories in our task suggest that the decision-making process can be completed after enough evidence has been accumulated (complete discrimination) or exited early before a complete decision has been made (incomplete discrimination). These clear effects on saccade trajectories can be applied to decision-making models, such as the recognition-primed decision model (Klein, 1999), recognition/metacognition model (Cohen et al., 1996), and OODA loop (Boyd, 2018), where quick decisions are made due to time constraints. The overall shape (amount and direction of saccade curvature) acts as a measure of how far along in the decision-making process the participant was at the point of saccade initiation (the exit point). Switching from manual to saccade responses in decision-making tasks can result in the ability to determine the stage at which the decision-making process was exited (early or after a complete decision is reached) and distinguish between these phases on a trial-by-trial basis.

### Conclusion

We varied distance and similarity in a saccadic response, visual search task using novel, complex objects. We found that the saccade metrics distinguished between active and complete decision-making processes, where distractor inhibition affected saccade curvatures sooner than saccade vector angles, suggesting that these are independent processes with separate priority maps. Distance had a strong influence on saccade trajectories in both stages, but the effect of similarity was limited to the active discrimination decision-making process. During this active stage, the effects of distance on saccade metrics followed spatial averaging and incomplete winner-take-all. When discrimination completed and the distractor was inhibited, a spatial suppressive surround mediated the effects of distance on saccade vector angles and curvatures. In comparison, an object-space suppressive surround mediated target-distractor similarity during the active discrimination process. When the distractor was later inhibited, decreasing similarity linearly increased the deviation of initial angle away from the distractor. We did not find any interaction between spatial and complex object discriminations, suggesting that distance and similarity are processed separately, consistent with spatial processing in the dorsal stream and object processing in the ventral stream. Distance and similarity differences must then independently feed into the oculomotor system for final saccade plans to be created. These results suggest that saccade responses would be more beneficial than manual responses in decision-making studies to allow for determining at what point in the decision-making process a decision was made based on these saccade trajectory metrics.

